# Influence of colour vision on attention to, and impression of, complex aesthetic images

**DOI:** 10.1101/2023.06.14.544891

**Authors:** Chihiro Hiramatsu, Tatsuhiko Takashima, Hiroaki Sakaguchi, Xu Chen, Satohiro Tajima, Takeharu Seno, Shoji Kawamura

## Abstract

Humans exhibit colour vision variations due to genetic polymorphisms, with trichromacy being the most common, while some people are classified as dichromats. Whether genetic differences in colour vision affect the way of viewing complex images remains unknown. Here, we investigated how people with different colour vision focused their gaze on aesthetic paintings by eye-tracking while freely viewing digital rendering of paintings and assessed individual impressions through a decomposition analysis of adjective ratings for the images. Gaze concentrated areas among trichromats were more highly correlated than those among dichromats. However, compared to the brief dichromatic experience with the simulated images, there was little effect of innate colour vision differences. These results indicate that chromatic information is instructive as a cue for guiding attention, whereas the impression of each person is unaffected by colour-vision genetics and would be normalised to their own sensory experience through one’s own colour space.

## 1. Introduction

Visual impressions are not mere representations of the external world captured by the retina.; they are subjective experiences produced by the workings of the entire visual system. The visual system, involving an active coordination between the eye and the brain, selects and concentrates on a limited portion of the image according to the viewer’s interests. The visual awareness gained through this active process results in subjective impressions unique to the individual, accompanied by emotional and aesthetic sensations [1–4]. Colour is one of the important factors influencing the formation of visual impressions [5–10]. Since colour perception is subjective, there is no guarantee that one person’s impression will be the same as another’s when viewing the same scene. Both inborn and acquired factors, such as culture, sex, and age, affect colour impression and preference [11–13].

Humans exhibit marked diversity in colour vision owing to differences in genetic background [14–16]. Although most humans have trichromatic colour vision, which is associated with three types of cones (S, M, and L), more than 200 million individuals have other colour vision types. Owing to polymorphisms in the L/M opsin genes encoded on the X chromosome, approximately 8% and 0.4% of European Caucasian men and women, respectively, and 4–5% and 0.2% of Asian (and presumably African) men and women, respectively, have these colour vision types [17–19]. These variations in colour vision mainly consist of trichromatic vision with closer spectral sensitivity between M and L cones (anomalous trichromacy) and dichromatic vision with only S and M (protanopia) or S and L cones (deuteranopia) [18].

Differences in colour discrimination ability among individuals with different types of colour vision have been well studied in the past [20–22]. Computer simulations have made it possible to create images that simulate the perception of dichromacy, and trichromats can now experience the expected perception of dichromats [23,24]. People with dichromatic vision who have S cones along with either M or L cones are generally thought to have difficulty discriminating colours on the red–green colour axis [25]. However, people with dichromatic vision responded similarly to people with trichromacy in colour-naming or categorical tasks if they experience a stimulus with sufficient magnitude and duration [26–29].

The impression of a scene viewed by a person who has experienced dichromatic or anomalous trichromatic vision for an extended duration may differ from that of a simulated image of those colours viewed by a person with trichromatic vision. People with dichromatic vision have sometimes similar semantic impressions of simple colours with people with trichromatic vision [30], despite differences in colour preference [31]. However, it is unclear whether this is also true for complex images that are rich in visual features. The human visual system has evolved to process various local visual features in the early visual cortex and integrate them into a unified perception in the higher visual cortex, which is influenced by experience during the course of development [32–34]. Therefore, individuals with different experiences may pay attention to complex images differently and have different impressions of them. In addition, it remains unknown how differences in colour vision affect the way people view complex visual scenes. Differences in colour vision may lead to differences in the saliency of complex images [35] and affect bottom-up visual attention. Attention to different parts of a complex image may subsequently lead to different impressions of the image (figure 1).

**Figure 1.**
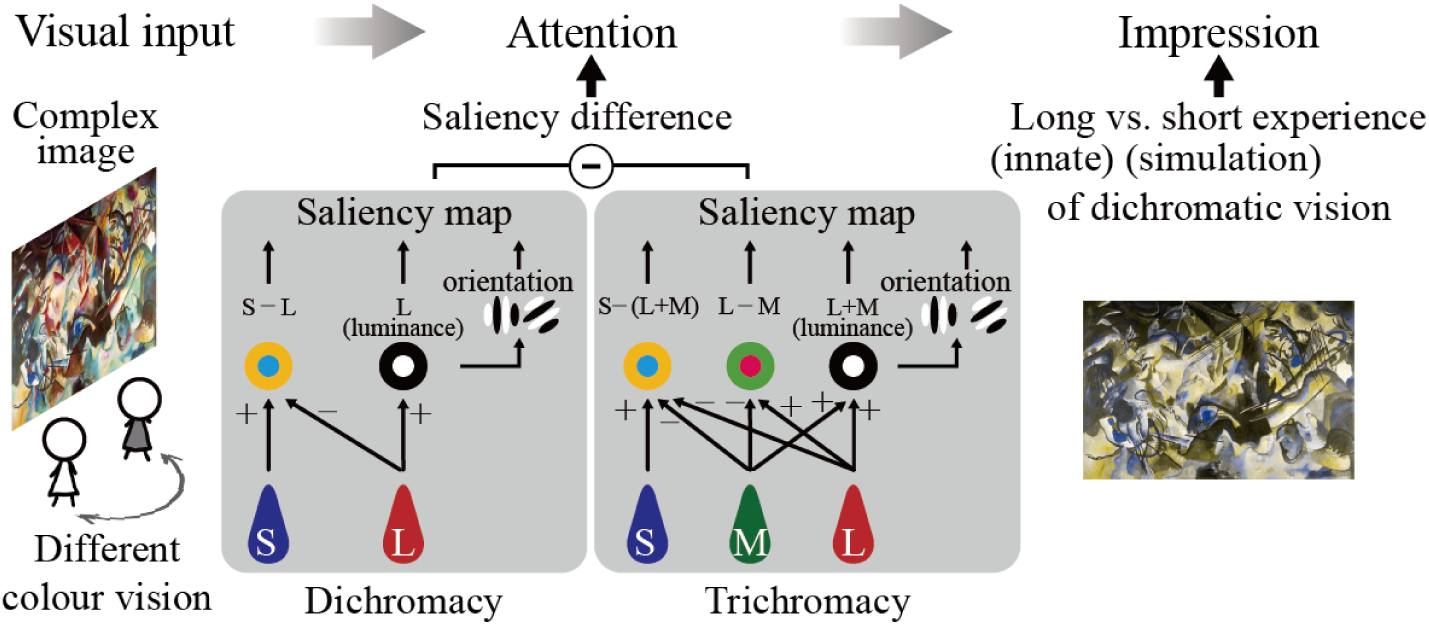
Schematic illustration of our study. The visual input of a complex image processed by the visual systems of individuals with different types of colour vision would result in differences in image saliency and attention [35], which may affect the viewer’s impressions of the image. The impression of the image may thus differ between people with a lifelong experience of congenital dichromatic vision and those who have briefly experienced stimulation-based dichromatic vision.

Although previous studies have extensively focused on how differences in colour vision affect abilities to discriminate, detect, name, or categorise colours [36–39], the diversity and commonality during the active process of viewing complex images remain largely unknown. Herein, we investigated the effect of congenital differences in colour vision on visual attention and viewers’ impressions of complex images. Using artistic paintings, in which a range of visual features (from low to high levels) have been shown to explain impressions, we extracted a variety of impressions, including colour and aesthetic sensations [7–10]. We devised an experiment wherein trichromatic observers were shown images that simulated dichromatic perception (hereafter, ‘simulated dichromats’) to compare the experience of congenital dichromacy with the short-term experience of simulated dichromacy.

## 2. Methods

### (a) Participants

Participants with diverse colour vision types were recruited through announcements on the website and in lectures. Written informed consent or assent (for underage participants) was obtained prior to the experiments. Participants were paid approximately $10 per hour. The participation criteria were as follows: sound mental health and uncorrected or corrected visual acuity sufficient to read the text on the screen. Fifty-eight participants (38 male, Mage = 30.3 ± 13.3 [SD]) from the community surrounding Kyushu University participated in the experiment. After predicting whether the participants had trichromacy, protan, or deutan on the Ishihara colour vision test plate, detailed colour vision type was determined via Rayleigh matching using an anomaloscope (Neitz OT-II, Neitz Co. Ltd., Tokyo, Japan). The Farnsworth–Munsell 100 Hue test (X-Rite, Grand Rapids, MI, USA) [40] was also conducted to resolve any discrepant findings. These colour vision tests are often used in colour vision research and clinical practice, and a battery of tests with the anomaloscope test increases reliability [21,41].

The L/M opsin genes of each participant were genotyped using long-range polymerase chain reaction (PCR) of *OPN1LW* and *OPN1MW*, and subsequent sequence analysis of three amino acid sites, 180, 277, and 285, which are important for the maximum absorption wavelength of M/L cone photoreceptor cells [42,43] (see details of genetic analysis below and table S2 for details regarding the participants’ colour vision).

One participant who reported a mental disorder was excluded from the analysis. Two underage participants and one participant whose first language was not Japanese were excluded from the impression analysis due to difficulty in interpreting adjectives. The gaze data of the two underage participants were excluded from the analysis to limit all data to adults. The number of participants and age range included in the gaze analysis for each colour vision group are as follows: those with dichromatic vision included 10 male participants (three with protanopic vision, seven with deuteranopic vision); Mage = 35.2 ± 16.5 (SD) (Mage = 37.6 ± 16.3 [SD] when four anomalous trichromats were included). Those with trichromatic vision included 10 male and 10 female participants: Mage = 30.5 ± 13.5 (SD). Trichromats who viewed dichromatic simulation images (simulated dichromats) included 11 male and 10 female participants: Mage = 26.1 ± 7.3 (SD). The trichromats and simulated dichromats did not overlap. All individuals participated in the experiment once.

### (b) Stimuli

To elicit diverse impressions, we used 24 digital images of abstract and figurative paintings with various colour and spacial configurations created by historically important deceased painters of the late 18th to early 20th centuries. These images were selected from among 684 paintings introduced in visual art sources such as WikiArt.org and omitted paintings that were realistic depictions of people, such as portraits of people. For the image selection, we used MDS analysis [44] with 16 parameters in images (maximum, minimum, mean, and variance for each value of CIExyY and those for predicted differences in saliency between trichromacy and dichromacy calculated based on the model proposed by Tajima and Komine [35]). By obtaining the position of each image in the two-dimensional MDS space analysed using the 16 parameters, dividing the MDS space into 4 × 6, and selecting 24 images from each region, we were able to select paintings with various colour and spacial information, free from preconceived notions and hypotheses. Dichromatic (deuteranopia) simulation images were created using the simulation software, Vischeck (www.vischeck.com), as a plugin for ImageJ [45]. The chromaticity and luminance of the display RGB primaries were measured using a spectroradiometer (SR- LEDW-5N, Topcon Technohouse, Tokyo, Japan) to calculate the CIExyY values of the stimuli.

### (c) Procedure

The experiment was conducted in a dark room. Images were presented on an OLED display (SONY PVM- 2541A, 24.5 inches, 1920 × 1080 pixels, native mode, maximum luminance = 116 cd^2^), and participants were positioned with their faces approximately 70 cm from the display and freely viewed each image for 30 s. The viewing angle was approximately 42.3 × 24.4°. A Tobii Pro X2-60 (Tobii Technology AB, Danderyd, Sweden) system was used for gaze measurement at a sampling rate of 60 Hz. After a valid 5-point calibration was performed, gaze measurements were obtained while participants viewed the paintings. The presentation of the images, control of the gaze measurement device, and data acquisition were performed using a custom program in Psychtoolbox3, run in MATLAB (Mathworks, Natick, MA, USA). After viewing each painting, participants rated their impression of the painting using the semantic differential method [46]. Twenty-three adjective pairs were selected based on previous studies: ‘ugly – beautiful’, ‘dark – bright’, ‘blurry – clear’, ‘disharmonious – harmonious’, ‘pale – thick’, ‘blunt – sharp’, ‘simple – complex’, ‘light – heavy’, ‘shallow – deep’, ‘loose – tense’, ‘calm – intense’, ‘soft – hard’, ‘gentle – flashy’, ‘dull – vivid’, ‘cold – warm’, ‘gloomy – cheerful’, ‘delicate – bold’, ‘unstable – stable’, ‘low contrast – high contrast’, ‘static – dynamic’, ‘monotone – colourful’, ‘weak – powerful’, and ‘dislike – like’ [47–49]. The adjectives were presented in Japanese. Participants rated each adjective pair between 1 (complete agreement with the adjective on the left) and 7 (complete agreement with the adjective on the right) by pressing a numeric key. The order of the images was randomly changed for each participant. The order of the adjective pairs was always the same. Each image remained visible during the impression rating. Half of the randomly selected participants with trichromatic vision observed dichromatic simulation images. The remaining trichromats observed the original images.

### (d) Analysis

#### Genetic analysis

DNA samples were extracted from participants’ buccal cells using the Buccal-Prep Plus DNA Isolation Kit (Isohelix, UK) and were anonymised at the Kyushu University Medical Information Center.

The first and second opsin genes (*OPN1LW* and *OPN1MW1* in the case of trichromacy) consisting of six exons spanning approximately 15 kbp on the X chromosome were amplified by polymerase chain reaction using long-range PCR [50,51]. Two primer sets were used for the first and second genes, respectively, as described below. The forward primers locate upstream of exon 1, and the reverse primer common for the first and second genes locate 3’ untranslated region of exon 6.

Primers for the first gene:

OPN1-P1-forward-1: 5′-GAGGCGAGGCTACGGAGT-3′

OPN1-E6-reverse-1: 5′-GCAGTGAAAGCCTCTGTGACT-3′

or

OPN1-P1-forward-2: 5′-AAGCCAACAGCAGGATGTGCG-3′

OPN1-E6-reverse-2: 5′-GCAGTGAAAGCCTCTGTGACTT-3′

Primers for the second gene:

OPN1-P2-forward-1: 5′-TTAGTCAGGCTGGTCGGGAACT-3′

OPN1-E6-reverse-1: 5′-GCAGTGAAAGCCTCTGTGACT-3′

or

OPN1-P2-forward-2: 5′-AAAGCCTAACAATGTCCAGGG-3′

OPN1-E6-reverse-2: 5′-GCAGTGAAAGCCTCTGTGACTT-3′

PCR was performed using 1.25 U PrimeSTAR GXL DNA Polymerase (Takara, Japan) with 0.2 µM of forward and reverse primers, 200 µM each of dNTP mix, 2 µL of genomic DNA, and an appropriate amount of buffer and purified water. The two-step PCR cycle was carried out as follows: 30 times repetition of 10 s at 98°C and 5 min 20 s at 68°C after 5 min denaturation process at 98°C.

The better amplified PCR product of the two primer sets was used for sequencing. First, PCR products were electrophoresed on a 0.8% agarose gel, and the amplified target region was cut out and purified utilising the illustra GFX PCR DNA and Gel Band Purification Kit (Cytiva, UK). PCR for direct sequencing was then performed using the following M13 tail primer pairs to sequence exons 3 and 5, which have three important amino acid loci (180 for exon 3 and 277 and 285 for exon 5) that contribute to different absorption wavelengths in L/M cone photoreceptors [42,43].

Primers for exon 3

OPN1-E3-M13-Forward: 5’-TGTAAAACGACGGCCAGTCCTTTGCTTTGGCTCAAAGC-3’ OPN1-E3-M13-Reverse: 5’-CAGGAAACAGCTATGACCGACCCTGCCCACTCCATCTTGC-3’

Primers for exon 5

OPN1-E5-M13-Forward: 5’-TGTAAAACGACGGCCAGTTCCAACCCCCGACTCACTATC-3’ OPN1-E5-M13-Reverse: 5’-CAGGAAACAGCTATGACCACGGTATTTTGAGTGGGATCTGCT-3’

Sequences were then determined directly by the Sanger method on an ABI 3730 genetic analyser using the BigDye direct cycle sequencing kit (ThermoFisher Scientific, USA). Presence or absence, and genotype of the first and second genes were determined based on the combination of three amino acids.

#### Gaze analysis

Attention maps were created from raw eye movement data. The raw data were considered valid if the output value of the eye-tracker reliability, ranging from 0 (most reliable) to 4 (least reliable), was less than 2 in either the left or right eye. For the x-y coordinates of the eyes, the average value of both eyes was used if both eyes were valid, and the value of one eye was used if only one eye was valid. We first created individual attention maps by calculating probability density functions for each image [52]. The time window was set to 5 s, during which the probability density of gaze at each image location was calculated. To reduce the effects of individual differences in noise in eye movement measurements, a two-dimensional Gaussian filter with a full-width at half-maximum of 3° was applied. The probability densities across the entire image were standardised such that the sum of the entire probability densities was 1 and the size of the attention map was reduced to 1/10 of the pixel size of the image. The time window was then shifted every second to obtain attention maps for the entire time-course. In addition, attention maps were created for cumulative time windows from the start of viewing, in 1-s steps after 5 s from the start of viewing.

Group attention maps for trichromats, dichromats, and simulated dichromats were created by summing and standardising individual attention maps weighted by the proportion of data validity during a time window (figure 2*a*).

**Figure 2.**
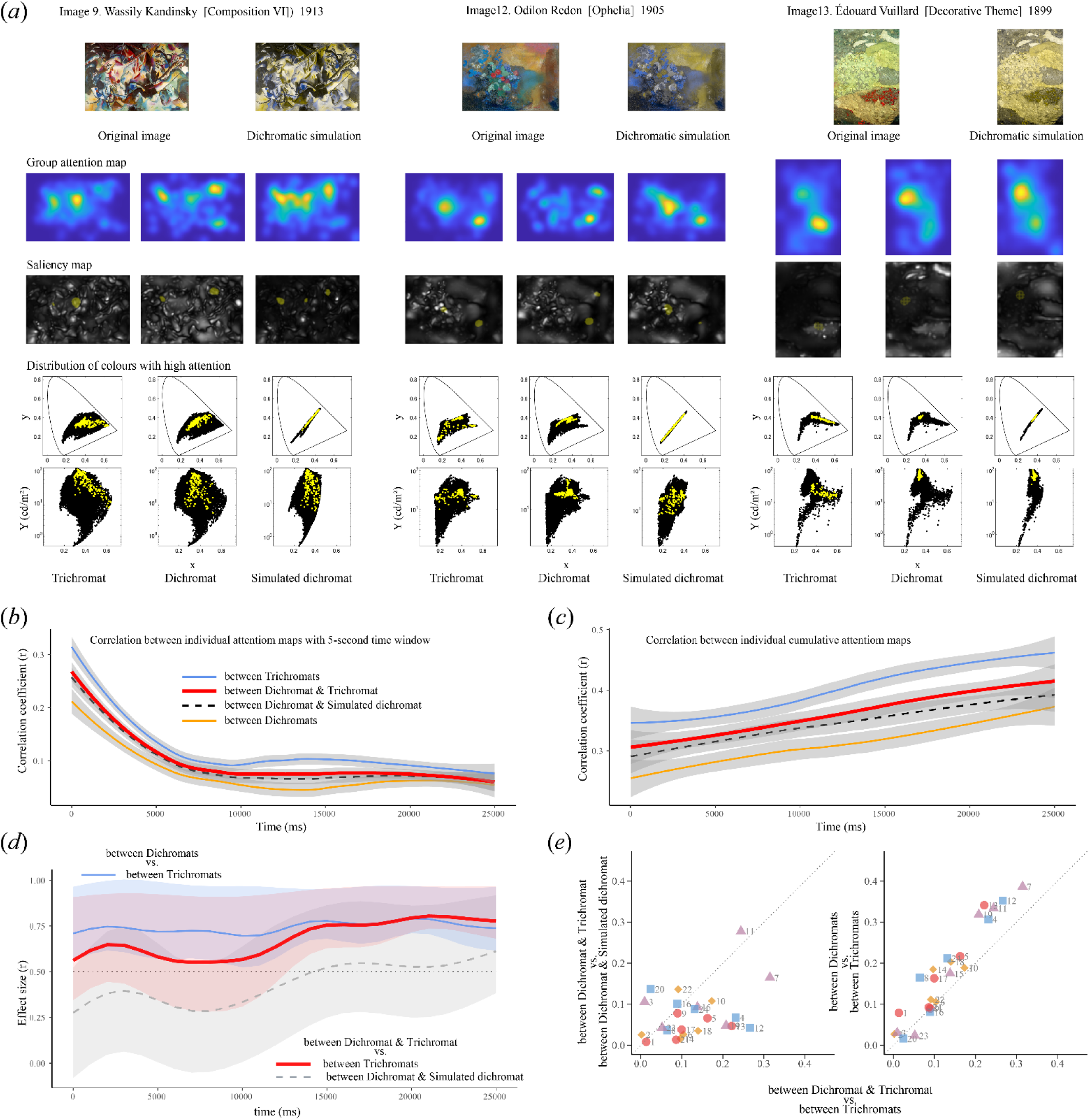
Gaze differences among colour vision types. (*a*) Examples of group attention maps for trichromats, dichromats, and simulated dichromats during the 0–5,000-ms observation period for images 9, 12, and 13. The original image viewed by trichromats and dichromats and the dichromatic simulation image viewed by half of the trichromats are shown above the attention maps. The saliency maps below the group attention maps are overlaid with the top 1% of pixels with high probability density in the group attention maps. The plots below the saliency maps show the distribution of image pixels in the CIE1931 xyY chromaticity diagram (top: x vs y, bottom: x vs Y; Y represents luminance). Black dots: distribution of all image pixels. Yellow dots: distribution of xyY values for the top 1% of the pixels with high attention. (*b*) Time-course of the pairwise correlations between individual attention maps with 5-s time window for various colour vision combinations. Each line indicates the smoothed mean value for medians of 24 images. Light blue line: trichromat and trichromat, bold red line: dichromat and trichromat, dashed black line: dichromat and simulated dichromat, orange line: dichromat and dichromat. Grey areas indicate 95% CIs. (*c*) Time-course of the pairwise correlations between individual attention maps with cumulative time window for various colour vision combinations. The symbols are the same as in panel *b*. (*d*) Time-course of the Wilcoxon effect size for the comparison of correlations between various colour vision combinations in *c*. Bold red line: dichromat and trichromat vs trichromat and trichromat; dashed grey line: dichromat and trichromat vs dichromat and simulated dichromat; light blue line: dichromat and dichromat vs trichromat and trichromat. Lines were smoothed with spline, and 95 % CIs are shaded. The dotted line represents r = 0.5. (*e*) Scatter plots of effect sizes of individual attention map differences for each image during 0–5,000 ms. Symbols beside image numbers indicate the categorisation of the images based on the values of the second dimension of the MDS for image selection. Red circle: extremely small; orange diamond: small; purple triangle: large; and light blue square: extremely large.

Pearson correlation coefficients between two individual attention maps for a given image were calculated for each time window for all participant combinations (pairwise correlation). For each image, the median of the correlation coefficients for the same colour vision combination was used as the representative value. The 24 medians from all images were used to illustrate the smooth mean line and 95 % CI for each colour vision combination (figure 2*b*, *c*) using the t-based approximation method called ‘loess’ in the ggplot2 library of R.

Saliency maps for each colour vision type were calculated based on Tajima and Komine’s model [35]. In the model, the local features in the luminance, chromatic, and orientation maps were extracted with centre-surround antagonism filters. Four spacial frequency scales and four orientation filters (vertical, horizontal, and two diagonal orientations) were used to extract orientation maps derived from the luminance signal. We assumed that L-M colour opponency was absent in the saliency map for deuteranopia. The weight on orientation maps relative to intensity/colour maps was set to 0.1. Saliency maps were obtained by normalising combined maps of orientation maps and intensity/colour maps. Saliency maps for simulated dichromats were obtained by using a deuteranopic simulation of the original image and estimating the saliency of a trichromatic viewer.

Using the median correlations between individual cumulative attention maps for the 24 images, the difference in correlation by colour vision combination was calculated as the Wilcoxon effect size and shown over time (figure 2*d*). The lower and upper bounds of the 95 % CIs of the effect sizes were estimated by bootstrapping with 1,000 replications. The Wilcoxon effect size was calculated using the correlation coefficients between the individual 0–5,000-ms attention maps for the differences in correlation between colour vision combinations in each image, and the magnitude of the differences in gaze between colour vision combinations was expressed as a scatter plot (figure 2*e*).

#### Impression analysis

To determine the effect of colour vision on individual impressions, the rating data for the 24 images based on 23 adjective pairs from 54 individuals were analysed using the Tucker decomposition method as described in the following equation [53,54].

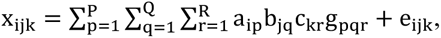

where *x_ijk_* is the three-phase data of the image *i ×* adjective *j ×* individual *k*. As in the above equation, the Tucker decomposition method represents *a_ip_* as the weighting of image *i* and latent factor *p*, *b_jq_* as the weighting of adjective *j* and latent factor *q*, and *c_kr_* as the weighting of individual *^k^* and latent factor *^r^*. *g_pqr_* is called the core tensor and reflects the importance of the interaction between factors. *e_ijk_* is the residual error. Each of the image phase, adjective phase, and individual phase was decomposed into three factors with orthogonality constraints. Therefore, *p*, *q* and *r* ranged from 1 to 3.

Differences in the weights of the three factors of individual phase among colour vision types were analysed using the Kruskal–Wallis test and the post-hoc Dunn test for multiple comparisons.

Gaze and impression data were analysed in MATLAB (version R2021b), whereas statistical analyses were performed in the R statistical environment (version 4-1.2, R Foundation, Vienna, Austria).

## 3. Results

### (a) Effect of colour vision on attention

The gaze analyses of genetically confirmed 10 dichromats, 20 trichromats, and 21 simulated dichromats during the 30-s free viewing of 24 painted images selected for having different image features revealed the diversity and commonality of attention to complex images and their time-course derived from differences in colour vision. Hereafter, ‘trichromats’ refer to individuals with normal cone absorption functions and exclude anomalous trichromats. In the main results section, we compared dichromats and trichromats, whereas the results, including four anomalous trichromats, are presented in the Supplementary Information.

#### Attention to example images

We investigated how participants with different colour vision types differed in terms of the areas to which they directed their attention when viewing complex images freely. figure 2*a* shows examples of the attention maps for the different colour vision groups created from individual attention maps that were calculated by applying the Gaussian function with a variance of 3° to the raw gaze data during the first 5 s of viewing. The saliency maps below the attention maps demonstrated the prediction of attention, which reflect the early bottom-up processes, including local spacial information for each colour vision type.

#### Differences in attention across colour vision types and images

Overall analysis of 24 images showed that the mean pairwise correlation coefficient reached a plateau at approximately r = 0.1 in attention maps, with 5-s time window at approximately 10 s (figure 2*b*), whereas it gradually increased until after 30 s in cumulative attention maps (figure 2*c*), regardless colour vision combinations.

The correlations between trichromats were consistently higher than those for other combinations, with the correlations between dichromats as the lowest; the correlations between dichromat and trichromat and those between dichromat and simulated dichromat were in between. Notably, the essential results did not change when anomalous trichromats were added to dichromats for analysis (figure S1) or when the analysis was limited to men to balance the number of participants.

The mean correlation coefficient between cumulative group attention maps and saliency maps for 24 images ranged around 0.1 for all colour vision types throughout the viewing period, although a slight improvement was observed for trichromat after 10 s (figure S2). Changing parameters, such as the number of spacial frequency scales or orientation filters and the weight of the orientation map in the saliency model, did not improve the overall correlation for the 24 images.

The time-course of the comparison of the individual cumulative attention map pairwise correlations between various colour vision combinations is shown in figure 2*d* as Wilcoxon effect sizes (r). The median values of the correlation coefficients between individual cumulative attention maps for the 24 images, which were used for the plots in figure 2*c*, were used to calculate the effect sizes and their 95 % CIs for the comparison of correlations. The Wilcoxon effect size for the comparison of correlations between dichromat and trichromat with correlations between trichromats (red bold line), and the comparison of correlations between dichromats with correlations between trichromats (light blue line) were greater than 0.5 throughout the time-course. Conversely, when correlations between dichromat and trichromat were compared with correlations between dichromat and a simulated dichromat (dashed grey line), effect sizes were lower than 0.5 for the first half of the viewing period.

To examine in which images the differences in colour vision affect the differences in attention map correlations, we obtained the effect size of the correlation comparison using the correlation coefficients between all individual attention maps during the first time window (0–5,000 ms) and represented the comparison between correlations in scatter plots (figure 2*e*). In the comparison of dichromat and trichromat correlations, and between trichromats, effect sizes were larger for images 5, 12, and 13 than for other images, while effect sizes were small for most images in the comparison of dichromat and trichromat correlations, and those between dichromat and simulated dichromat (figure 2*e* left). The colour of the symbols indicates the position in the multidimensional scaling (MDS) analysis [44] for the image selection process. The images indicated by red circles, which have extremely small values in the second dimension of the MDS with tendency of having red– green or lightness spacial contrast, are plotted below the diagonal line in the left plot of figure 2*e* except for image 21, indicating that those images garnered different attention from dichromats and trichromats. In the comparison of correlations between dichromats and between trichromats, effect sizes were also larger for the specific images, therefore each image is plotted near the diagonal line in figure 2*e* right. These results indicate that the attention maps were more similar among trichromats than among other colour vision combinations for particular images.

When anomalous trichromats were added to dichromats for these analyses, the results showed the same tendency, although gaze differences with trichromats were mitigated (figure S1).

### (b) Effect of colour vision on impression

To evaluate the effect of colour vision on the impressions of individuals with different colour vision types, the impression rating data (24 images × 23 adjective pairs × 54 individuals) were decomposed into three factors for each of the image, adjective, and individual phases using the Tucker method [53]. The decomposed model explained 90.8% of variation.

#### Individual factors

For factor 3 of the individual phase, the distributions of trichromats (open circles) and simulated dichromats (black circles) differed markedly, and the distribution of dichromats (triangles: protanopia, and inverted triangles: deuteranopia) and anomalous trichromats (diamonds) appeared to overlap with the distribution of trichromats. No differences were observed among the two types of dichromats (deuteranopia and protanopia) and anomalous trichromats in terms of factor 3 (χ^2^ = 1.14, *p* = 0.57, Kruskal–Wallis test). Therefore, dichromats with protanopic or deuteranopic vision were combined into a single group for analysis. There were differences among trichromats, dichromats, and simulated dichromats for factor 3 (χ^2^ = 23.34, *p* < 0.0001, Kruskal–Wallis test). A post-hoc Dunn test for multiple comparisons indicated that simulated dichromats differed significantly from dichromats (*p.adj* < 0.001) and trichromats (*p.adj* < 0.001), as shown in the lower panel of figure 3*b*. For factors 1 and 2, there were no significant differences among the colour vision groups (top and middle panels of figure 3*b* (factor 1: χ^2^ = 3.31, *p* = 0.19; factor 2: χ^2^ = 1.32, *p* = 0.52, Kruskal–Wallis test). The results were almost identical when dichromats and anomalous trichromats were analysed as a single group.

**Figure 3.**
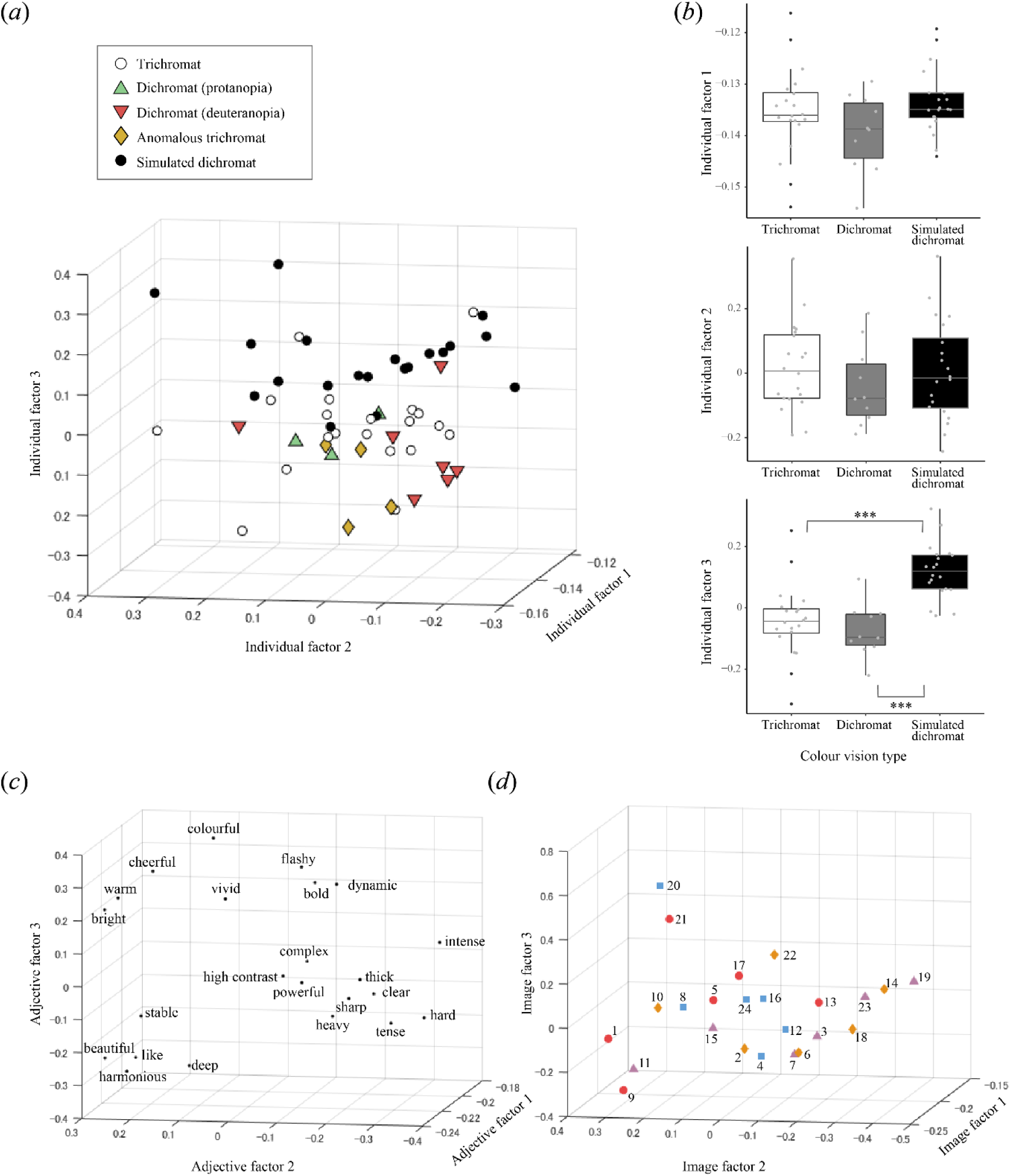
Effects of colour vision on impressions. (*a*) Distribution of weights for individuals with different colour vision types in a three-dimensional space consisting of individual factors 1–3. (*b*) Box plots and distribution of weights for each individual factor in trichromats, dichromats, and simulated dichromats. The horizontal line of each box indicates the median value. The box represents the range between the first and third quartiles. The upper and lower whiskers refer to the largest and smallest data points in the range from the first quartile − 1.5*the range of box to the third quartile + 1.5*the range of box, respectively. Each dot represents one data point. ***adjusted *p* < 0.001. (*c*) Distribution of weights for adjectives in a three-dimensional space consisting of adjective factors 1–3. (*d*) Distribution of weights for images in a three-dimensional space consisting of image factors 1–3. The symbols beside the image numbers indicate the categorisation of the images based on the values of the second dimension of the MDS for image selection. Red circle: extremely small; orange diamond: small; purple triangle: large; and light blue square: extremely large.

#### Relationship among factors in the image, adjective, and individual phases

The core tensor G of the Tucker decomposition (table 1) reflects the interactions among the three-phase factors. Factor 3 in the individual phase exhibited a large positive interaction with image factor 1 and adjective factor 3, as well as a large negative interaction with image factor 3 and adjective factor 1.

**Table 1.** Core tensor G in the Tucker decomposition

| Individual | Image | Adjective |  |  |
| --- | --- | --- | --- | --- |
|  |  | Factor 1 | Factor 2 | Factor 3 |
| Factor 1 | Factor 1 | −714.6921 | 0.1215 | −0.2572 |
|  | Factor 2 | 0.2714 | 96.3313 | 0.9473 |
|  | Factor 3 | −0.0334 | −0.0923 | 54.1160 |
| Factor 2 | Factor 1 | −5.0401 | −5.1168 | 8.5798 |
|  | Factor 2 | −3.7604 | −31.0129 | 0.1383 |
|  | Factor 3 | 6.0097 | −1.2950 | −11.2864 |
| Factor 3 | Factor 1 | 0.6185 | −3.0068 | 25.8936 |
|  | Factor 2 | 4.0170 | 1.7134 | 0.2417 |
|  | Factor 3 | −16.8498 | 4.6614 | 5.2207 |

The magnitude of the value reflects the importance of the interactions between the factors in the three phases.

The adjective ‘colourful’ had the largest positive value for adjective factor 3 (figure 3*c*). Multiplication of the weights of adjective factor 3 for colourful (0.3937), image factor 1 (all negative, figure 3*d*), individual factor 3, and the core tensor value (25.8936, table 1) that connects three-phase factors contributes the reconstructed rating value. Therefore, the more negative the weights of individual factor 3, the higher the reconstructed rating of ‘colourful’. The interaction among individual factor 3, image factor 3, and adjective factor 1 was negative (−16.8498, table 1). Since all weights of adjective factor 1 were negative, more negative image factor 3 and individual factor 3 values yielded a higher impression.

Thus, among simulated dichromats, a high weight of individual factor 3 indicated a reduced colour impression. This was consistent with the distribution of raw rating data for simulated dichromats (figure S3).

figure S4*a* shows the average impression ratings for the colour vision groups, ordered by the lowest weight of image factor 3 and highest weight of adjective factor 3. Evaluating the difference between groups highlights which images and adjectives are more affected by colour vision conditions (figure S4*b*). When dichromats and simulated dichromats were compared, we observed large differences in images with low image factor 3 weights, particularly for adjectives with high adjective factor 3 weights such as ‘colourful’ and ‘flashy’. This phenomenon was also observed for comparisons between trichromats and simulated dichromats. However, no systematic differences were observed between dichromats and trichromats.

#### Impression analysis with restricted images

Finally, we examined whether the impression analysis used in this study is a valid method for detecting innate colour vision differences. A posteriori, we restricted impression data to images in which differences between dichromats and trichromats were prominent for some adjectives, as shown in figure S4*b*. For example, we performed Tucker decomposition using impression data from images 1 and 12 and found that innate colour vision differences appeared in individual factors 1 and 3 (figure S5). In this case, the decomposed model explained 92.5% of the variability. The values of the image factor were all negative. The values of individual and adjective factors in figure S5, and core tensor G in table S1 indicate that the images were seen more colourful or complex as smaller the individual factor 1 was, or more bold, dynamic, and cheerful as larger the individual factor 3 was. Although we cannot deduce the conclusion from this result alone because of the limited number of images, it indicates that the methods we used in this study were valid to reveal the factors such as differences in colour vision that influence impression formation.

#### Other factors influencing impression

Additional multiple regression analyses using colour vision type (trichromats, simulated dichromats, dichromats, and anomalous trichromats), age, sex, and the 100 Hue Test error score, as explanatory variables, showed that none of these factors, except the colour vision type (*F* = 8.03, *p* < 0.001), influenced individual factor 3. Age had a significant effect on individual factor 2 (*F* = 4.24, *p* < 0.05), which exhibited a large negative interaction (−31.0129, table 1) with adjective factor 2 and image factor 2 (figure 3*c*, *d*).

## 4. Discussion

Gaze analyses revealed that trichromats have higher attentional similarity than dichromats. In contrast, impression analyses indicated an overall similarity between dichromats and trichromats for paintings with varying visual features, at least in subjective evaluation. Overall, the attention map correlations for the various images revealed that correlations between trichromats were always higher than those of the other combinations, whereas correlations between dichromats were always lower. Although the data were not shown (to avoid complexity), the correlation between simulated dichromats during 0–5,000 ms was nearly identical to that between dichromats and trichromats. These results suggest that images with no red–green colour information may result to large gaze variability, likely reflecting the differences in the bottom-up information that individuals place weight on. The individual differences among trichromats became smaller as the amount of chromatic information increased. This may indicate one of the significances of the red–green axis in trichromatic vision.

We found that the correlation between dichromats and simulated dichromats was not as high as those between trichromats and that the effect size of differences from the correlation between dichromat and trichromat was low. This may be attributed to the large individual differences between the attention maps of dichromats and simulated dichromats, where some dichromats were more similar to trichromats, while some dichromats were more similar to simulated dichromats.

In several images, colour vision affected the attention directed toward the images within the first 5,000 ms of viewing with larger effect size. One example of an obvious difference between colour vision types is that for image 13, in which both red–green and lightness contrast occupy space, trichromats directed their attention to the dark red locations at the bottom of the image, whereas dichromats directed their attention to the light area in the upper left corner of the image. The distribution of the pixels with high gaze probability density in the CIExyY plots clearly shows this tendency.

The tendency of colour and spacial composition of the images that engendered large differences in gaze between colour vision types can be estimated from the MDS values that were used for experimental image selection and classification. In images with high red–green or lightness contrasts and extremely small values in the second dimension of the MDS, attention maps differed between dichromats and trichromats, whereas in images with high blue–yellow or unsaturated colours, red–green colour vision had little effect. For these images, the correlations of the attention maps were low owing to high individual variation, even among participants with the same colour vision type, or the correlations were high, regardless of colour vision type. However, image 12 was an exception, as it demonstrated higher attention map correlations among trichromats than other colour vision combinations despite having high blue–yellow contrast. This was presumably due to the red floral objects near the centre of the image. Overall, these results suggest that colour vision influences attention to an image in the early stages of viewing (approximately within 5,000 ms), depending on the spacial colour distribution of the image.

The difference in dichromatic and trichromatic saliency was used as one of the parameters of the MDS analysis for image selection. These facts indicate that differences in saliency, including spacial information, are useful in predicting the impact of innate differences in colour vision on bottom-up attention to complex images. However, overall low correlations between saliency maps and attention maps, even in trichromats, suggest that the saliency model based solely on bottom-up processes is insufficient to predict viewer attention, which involves top-down attention to visual contents and gaze centre bias [55,56]. The low correlation of the attention maps among dichromats may suggest that dichromats’ weighting of bottom-up information varies. Therefore, it is important to recognise that having a univocal interpretation is challenging, especially when estimating how people with dichromatic vision view the world.

The difference in impression among the participants is essentially driven by the fact that trichromats viewing images made to simulate the experience of a dichromat do not find those images colourful. Meanwhile, no obvious differences were observed between the impressions of congenital dichromats and trichromats when 24 images containing various visual features were analysed. These results indicate that short-term experiences of dichromatic vision using simulation images have very different effects on visual impressions than innate colour vision.

The results of the impression analysis may reflect the relative assessments of the observers, where participants rated adjectives relative to the range of stimuli they have experienced. According to this interpretation, simulated dichromats may differ from dichromats because the colour losses in the simulated images were apparent to them, although most simulated dichromats were unaware that they were observing simulated images until after the experiment. Nonetheless, congenital dichromats experienced a rich impression, although it was relative to each observer. A previous study that examined the influence of lens-brunescence on colour naming found that the results from elderly colour vision simulations was not consistent with those from intrinsic colour vision of older observers, suggesting the influence of lifelong chromatic adaptation [57].

These observations are consistent with the recent view that the plasticity of post-receptoral processes calibrates variations between individuals during experience-dependent development to discount congenital variations of receptors [58], although plasticity sometimes brings about striking individual differences in colour perception, as revealed by the image #thedress [59]. The possibility of compensating for red–green contrast loss in anomalous trichromats at post-receptoral stages has been investigated by performing psychophysical experiments [60,61]. Physiological studies using simple colour stimuli have shown that signals are amplified in the brains of people with anomalous trichromatic vision [62,63]. These compensations may occur when they view complex images. It is also plausible that dichromats compensate similarly by using additional information obtained from rod or melanopsin mechanisms and variations in the optical density of cones, macular pigment, lens, among others [22,64–66].

Although the colour space that an individual can experience is limited by genetic factors, our findings suggest that individuals with different colour vision can optimise the information available in the intrinsic colour space and construct impressions, including colour impressions, for various complex scenes. The colour impressions constructed in this way may yield impression strength of the same level when viewed in relation with other impressions, even if the colour space is different.

The influence of congenital colour vision on impressions cannot be completely ruled out, although it was not pronounced across the various images used in this study. The raw rating data for ‘sharp’ was greater in dichromats and anomalous trichromats than in trichromats. Furthermore, when impression data were restricted on certain images a posteriori, individual factors 1 and 3 of Tucker decomposition differed between trichromats and other congenital colour vision types. People with dichromatic or anomalous trichromatic vision may have a spectacular impression of an image that has a clear colour-luminance contrast for them. Image 12 used in this additional analysis obtained a higher rating for ‘colourful’ among dichromats than among trichromats (figure S4*b*); attention analysis showed that this image had a higher similarity in attention maps among trichromats than among dichromats. Since differences in attentional positions were mainly reflection of the attention in the early stage of viewing, differences in impressions may be more pronounced for shorter viewing times. Further studies are required to investigate these possibilities. In addition, the comparison between dichromats and simulated dichromats demonstrated differences in a relatively large number of adjective pairs (figure S3), further indicating that it is difficult to infer the impression of dichromats by using simulation images alone.

Overall, our results indicate that even when people with trichromatic vision receive the attenuated colour impression for dichromacy-simulating images, their subjective experience differs from that of someone who has experienced dichromatic vision throughout life. This may be of relevance because there is a lot of cerebral cortex within so-called ‘high-level’ visual areas involved in colour, and these areas do not mature until years after birth [34]. Colour vision is an active process that connects sensory signals with meaning; considering the vast cortical areas processing colour information, it is plausible that the brain is not only capable but has also evolved to collect sufficient information from the retina (even in dichromats) to extract meaning from retinal images.

Additional multiple regression analysis showed that age also affected the individual difference in impression. As age increased, individual factor 2 decreased, suggesting fewer extreme evaluations of impressions, such as ‘beautiful’, ‘harmonious’, and ‘bright’. Studies with larger sample sizes from various ages should further clarify this. Since the error score on the 100 Hue Test did not affect the impression, our findings suggest that differences in colour discrimination have little effect on impression.

This study has a few limitations. First, owing to the limited number of dichromats and anomalous trichromats, they participated only in the experimental condition wherein they viewed original images. Some dichromats viewed the dichromatic simulation images after the experiment and reported that they appeared similar to the original images. Pioneering simulation studies on dichromacy also reported that dichromats were satisfied with the agreement between the simulated image and the original image [23,24]. Therefore, if dichromats had evaluated their impressions upon viewing the dichromatic simulation images during the experiment, the results would likely have been similar. As we used deuteranopic simulation as a representation of dichromatic simulation, three protanopic dichromats might be able to discriminate between original and simulated images if compared. Nevertheless, the threshold for red–green contrast was diverse and continuous in dichromats and anomalous trichromats [67]; therefore, a single transformation cannot precisely simulate the perception of dichromats or anomalous trichromats. Whether subtle differences affect the attention to and impression of complex images requires further investigation. Second, although we used paintings of high artistic qualities, which are suitable for obtaining low-to high-order semantic evaluations, it would be important to examine how differences in colour vision are affected when observing natural images. Third, this study did not explain how differences in lower-order visual features are integrated to produce a similar level of higher-order impressions. These limitations should be addressed in future studies. Examining brain activity reflecting visual, emotional, and cognitive processes and personal preferences may improve our understanding of the mechanisms underlying the formation of impressions observed here.

## 5. Conclusion

We investigated how differences in colour vision types affected attention to and impressions of complex images, which encompassed paintings consisting of various colour configurations. Innate colour vision was found to influence attention in the early stages of image viewing. Meanwhile, the colour impressions of those who had a temporary dichromatic experience generated using dichromatic simulation images decreased, whereas there were no systematic differences in impressions between individuals with congenital dichromatic vision and those with trichromatic vision. Thus, dichromatic simulation images, which are theoretically generated based on the colour discrimination ability of dichromacy, can be a predictor of behaviours influenced by bottom-up attention to some extent but cannot infer the impression experienced by dichromats. Human colour vision exhibits significant diversities beyond the classifications used in this study, including variations in genetic factors [16,18,66,68,69,67,70]. Little is known regarding the plasticity and compensation mechanisms involved in post-receptoral processes [22,58,71]. Exploring these unknowns and examining how the diversity of colour vision affects attention, impressions, and emotions will provide a better understanding of the actual effects of diversity on behaviour and subjectivity.

## Ethics

This study was approved by the Ethics Committee for Human Experiments (Approval No. 241) and the Ethics Review Committee for Human Genome and Genetic Analysis Research (Approval No. 651) of Kyushu University, Japan.

## Data accessibility

The gaze and impression data and codes for the main analyses have been deposited in the DRYAD repository created for this project (doi:10.5061/dryad.w6m905qs5).

## Authors’ contributions

C.H. and S.T. designed the study; C.H., T.T., H.S., and X.C. performed the research; C.H., T.T., and H.S. analysed the data; S.T. provided codes for saliency analysis; T.S. and S. K. provided experimental equipment; and C.H., T.S., and S.K. discussed the results and wrote the paper.

## Acknowledgements

We thank Hidenori Tachida and Etsuko Moritsuka for providing the genetic analysis environment, Ryuichi Ashino and Yuka Matushita for supporting the genetic analysis, Hisao Ueyama for providing the latest protocol for long-range PCR, Hiroaki Oie for helping to recruit the participants. We dedicate this paper as a tribute to the memory of our collaborator, Satohiro Tajima, who passed away in 2017.

