## Supplementary Information for "Influence of colour vision on attention to, and impression of, complex aesthetic images"

**This PDF file includes:**

Figures S1 to S5
Tables S1 to S2

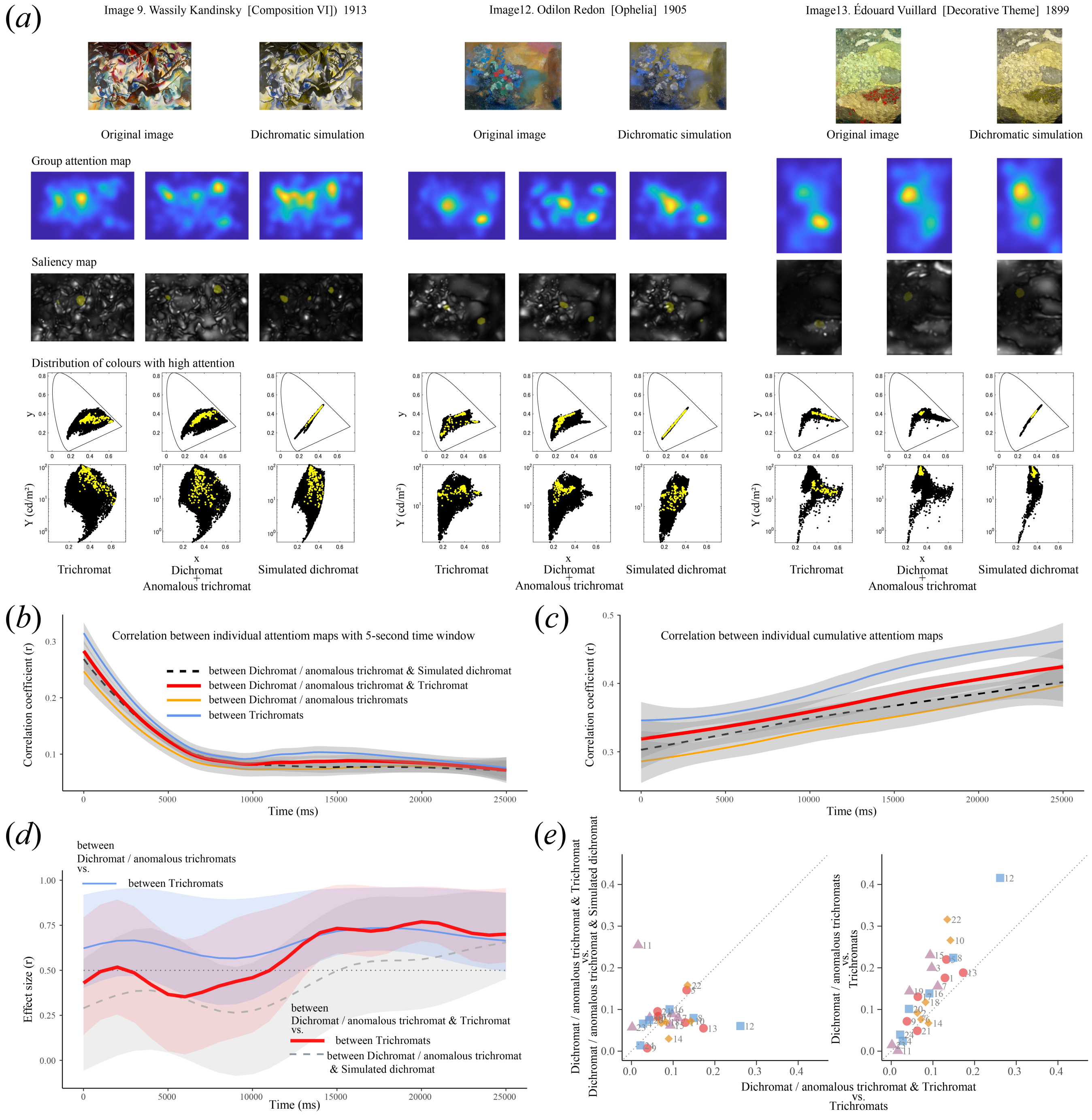


**Figure S1.** Gaze differences among colour vision types. (*a*) Examples of group attention maps for trichromats, dichromats and anomalous trichromats, and simulated dichromats during the 0–5,000 ms observation period for images 9, 12, and 13. The original image viewed by trichromats and dichromats/anomalous trichromat as well as dichromatic simulation image viewed by half of the trichromats are shown above the attention maps. The saliency maps below the group attention maps are overlaid with the top 1% of pixels with high probability density in the group attention maps. The plots below the saliency maps show the distribution of image pixels in the CIE1931 xyY chromaticity diagram (top: x vs. y, bottom: x vs. Y; Y represents luminance). Black dots: distribution of all image pixels. Yellow dots: distribution of xyY values for the top 1% of the pixels with high attention. (*b*) Time-course of the pairwise correlations between individual attention maps with 5-s time window for various colour vision combinations. Each line indicates the smoothed mean value for medians of 24 images. Light blue line: trichromat and trichromat, bold red line: dichromat/anomalous trichromat and trichromat, dashed black line: dichromat/anomalous trichromat and simulated dichromat, orange line: dichromat/anomalous trichromat and dichromat/anomalous trichromat. Grey areas indicate 95% CIs. (*c*) Time-course of the pairwise correlations between individual attention maps with cumulative time window for various colour vision combinations. The symbols are the same as in panel *b*. (*d*) Time-course of the Wilcoxon effect size for the comparison of correlations between various colour vision combinations in *c*. Bold red line: dichromat/anomalous trichromat and trichromat vs trichromat and trichromat; dashed grey line: dichromat/anomalous trichromat and trichromat vs dichromat/anomalous trichromat and simulated dichromat; Light blue line: dichromat/anomalous trichromat and dichromat/anomalous trichromat vs trichromat and trichromat. Lines were smoothed with spline, and 95% CIs are shaded. The dotted line represents r = 0.5. (*e*) Scatter plots of effect sizes of individual attention map differences for each image during the 0–5,000 ms. Symbols beside image numbers indicate the categorisation of the images based on the values of the second dimension of the MDS for image selection. Red circle: extremely small; orange diamond: small; purple triangle: large; and light blue square: extremely large.

(*a*)

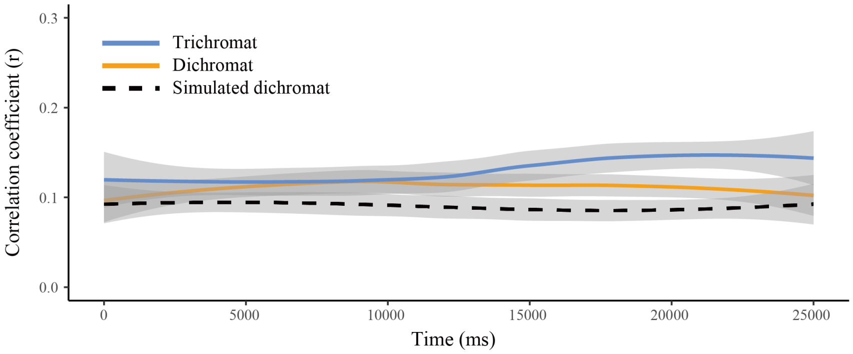


(*b*)

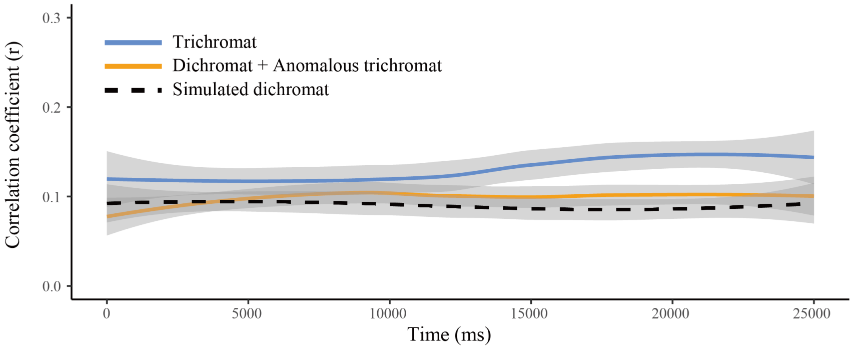


**Figure S2.** Time-course of the correlations between group cumulative attention maps and saliency maps. (*a*) Pearson correlation coefficients obtained from the 24 images for each time window were used to plot the smooth mean line and 95% CI for each colour vision type by the "loess" method. Light blue line: trichromat; orange: dichromat; dashed black line: simulated dichromat. Grey areas indicate 95% CIs. The same saliency map was used for all time windows. (*b*) Same analysis in A, including anomalous trichromats.

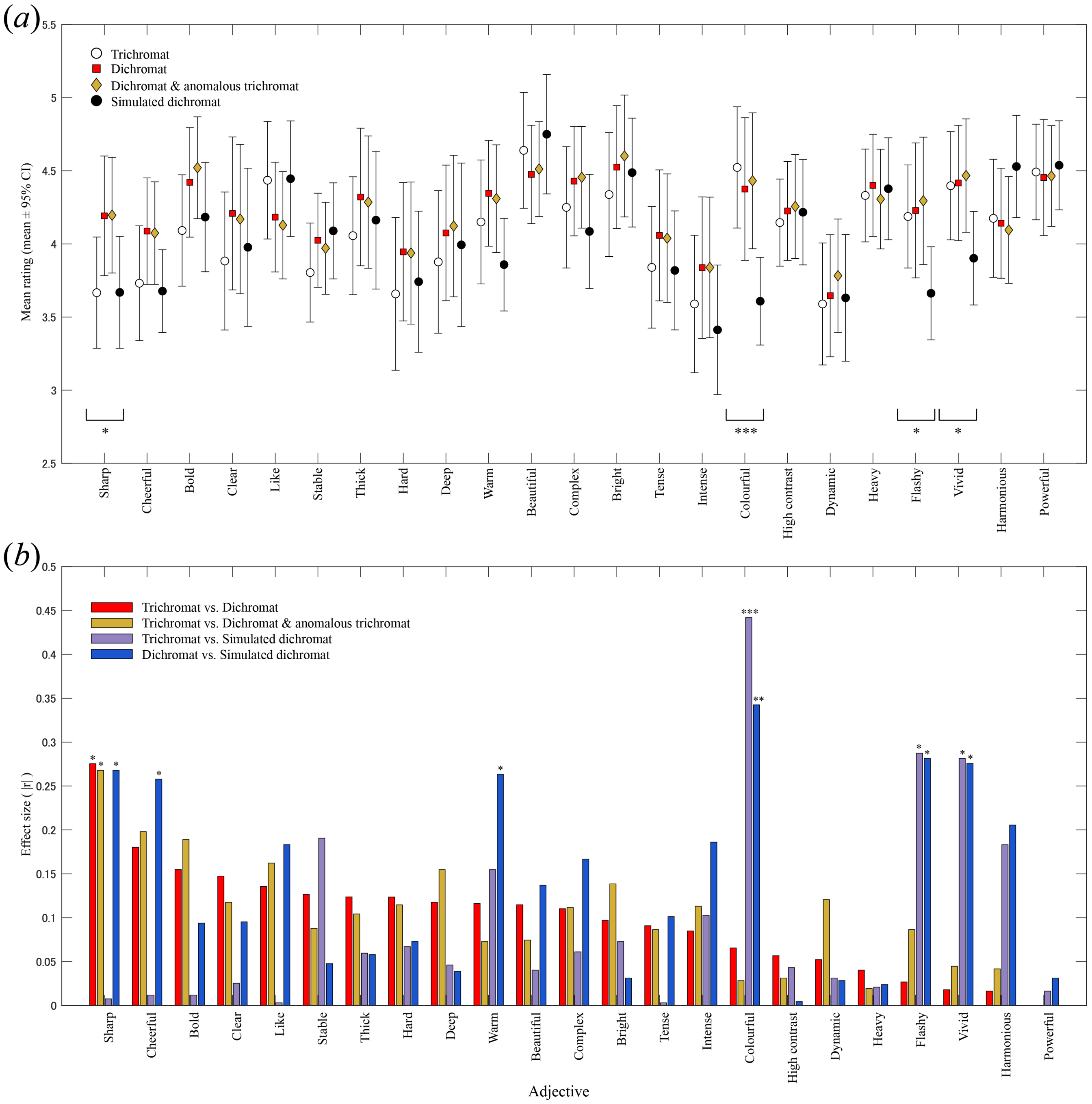


**Figure S3.** Rating differences for 23 adjective pairs among colour vision types. (*a*) Mean ± 95% CI of ratings for 24 images for each colour vision type. Open circle: trichromats; red square: dichromats; orange diamond: dichromats and anomalous trichromats; and black circle: simulated dichromats. Labels indicate adjectives that were assigned a larger value. *** $p < 0.01$, * $p < 0.1$ (marginally significant) in Kruskal Wallis test. (*b*) Mann-Whitney effect size (absolute r) for comparison between each colour vision type. Red: trichromat vs dichromat; orange: trichromat vs dichromat and anomalous trichromat; purple: trichromat vs simulated dichromat; and blue: dichromat vs simulated dichromat. Adjective pairs are shown in descending order based on the absolute values of Mann-Whitney effect size between trichromat and dichromat. Asterisks indicate the significance of differences by uncorrected p-values. *** $p < 0.01$, ** $p < 0.05$, * $p < 0.1$.

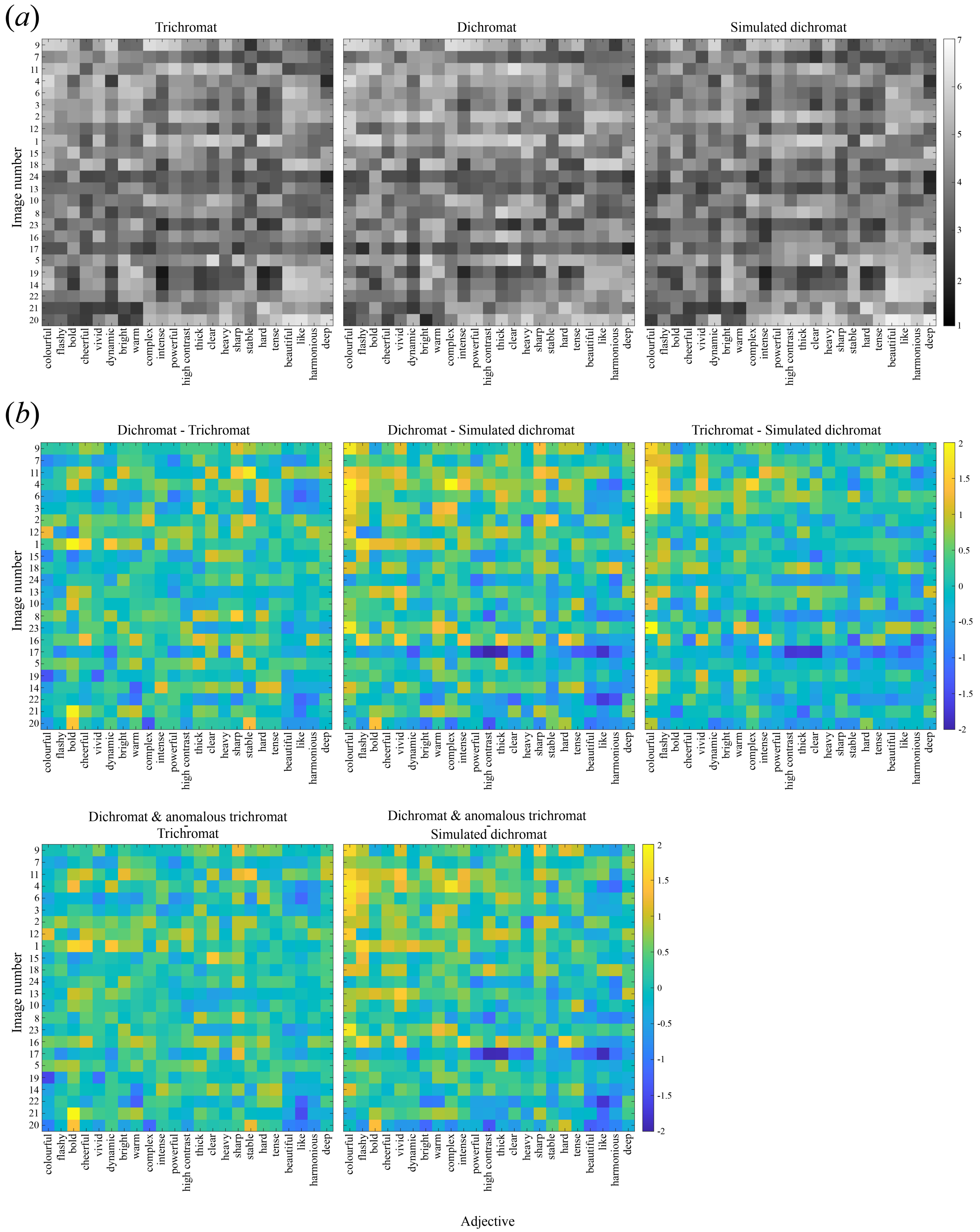


**Figure S4.** Adjective ratings for each image. (*a*) Grey scale indicates the average rating in each colour vision group. (*b*) Colour scales indicate differences between colour vision groups. Contrasts between dichromat and anomalous trichromat vs. trichromat are also shown for reference. Images are arranged in ascending order of the weight of image factor 3 in the Tucker decomposition, and adjectives are arranged in descending order of the weight of adjective factor 3.

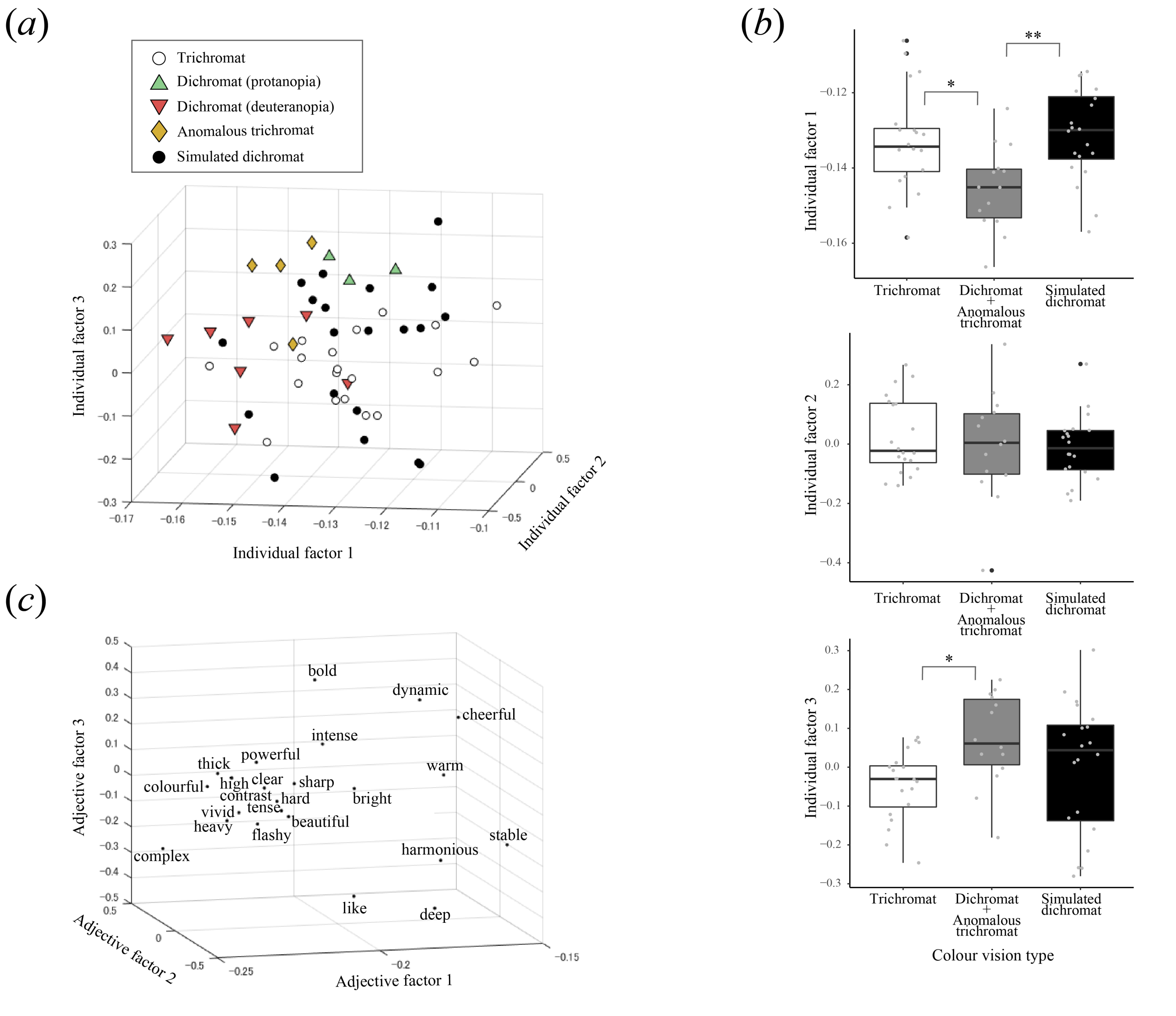


**Figure S5.** Effects of colour vision on impressions for restricted images (images 1 and 12). (*a*) Distribution of weights for individuals with different colour vision types in a three-dimensional space consisting of individual factors 1–3. (*b*) Box plots and distribution of weights for each individual factor in trichromats, dichromats and anomalous trichromats, and simulated dichromats. The box represents the range between the first and third quartiles. The upper and lower whiskers refer to the largest and smallest data points in the range from the first quartile - 1.5*the range of box to the third quartile + 1.5*the range of box, respectively. Each dot represents one data point. ** $p.adj < 0.01$, * $p.adj < 0.05$. (*c*) Distribution of weights for adjectives in a three-dimensional space consisting of adjective factors 1–3.

**Table S1. Core tensor G in the Tucker decomposition with restricted images**

| | Adjective factor 1 | Adjective factor 2 | Adjective factor 3 |
|---|---|---|---|
| Individual factor 1 | -218.1786 | 0 | 0 |
| Individual factor 2 | 0 | -18.5425 | 0 |
| Individual factor 3 | 0 | 0 | -14.3383 |

The magnitude of the values reflects the importance of the interactions between the factors in the three phases. Because of the small number of images, the number of factor for the image phase was set to 1, while the individual and adjective phases were set to 3.

**Table S2. Detailed colour vision information of participants with dichromatic or anomalous trichromatic vision**

| Anonymized participant code | Anomaloscope result | Anomaloscope anomalous quotient | 100 Hue test error score | *OPN1LW* (1st gene) a.a. 180-277-285 | *OPN1MW* (2nd gene) a.a. 180-277-285 |
|---|---|---|---|---|---|
| 1 | protanopia | $\infty$ - 0 | 56 | A-F-A | A-F-A |
| 2 | protanopia | $\infty$ - 0 | 72 | A-F-A | A-F-A |
| 3 | protanopia | $\infty$ - 0 | 72 | A-F-A | A-F-A |
| 4 | deuteranopia | $\infty$ - 0 | 92 | A-Y-T | A-F-A |
| 5 | deuteranopia | $\infty$ - 0 | 136 | S-Y-T | A-F-A |
| 6 | deuteranopia | $\infty$ - 0 | 256 | S-Y-T | A-F-A |
| 7 | deuteranopia | $\infty$ - 0 | 168 | S-Y-T | not amplified |
| 8 | deuteranopia | $\infty$ - 0 | 204 | A-Y-T | N-F-A |
| 9 | deuteranopia | $\infty$ - 0 | 256 | S-F-A | S/A-F-A |
| 10 | deuteranopia | $\infty$ - 0 | 148 | not amplified | not amplified |
| 11 | deuteranomaly | $\infty$ - 0.26 | 176 | A-Y-T | not amplified |
| 12 | deuteranomaly | 7.64 - 3.21 | 160 | S-Y-T | S-Y-T |
| 13 | deuteranomaly | 7.64 - 2.33 | 40 | not amplified | not amplified |
| 14 | deuteranomaly | 4.69 - 2.33 | 20 | A-Y-T | A-Y/F-T/A |

The results of the colour vision test using the anomaloscope, the anomalous quotient, the error score of the 100 Hue test, and the combination of amino acid sites 180, 277, and 285 in the opsin proteins coded by the first and second L/M opsin genes (*OPN1LW* and *OPN1MW*) are listed. A range of $\infty$ to 0 in anomalous quotient indicates dichromacy, and an interval in between $\infty$ to 0 indicates anomalous trichromacy. Amino acids are indicated by one-letter abbreviations; A: alanine, F: phenylalanine, S: serine, T: threonine, and Y: tyrosine. The three-site combination of the first gene in trichromacy is S-Y-T or A-Y-T, and the three-site combination of the second gene is usually A-F-A. Among trichromats, the 31% of the X chromosome had A at the 180th site of the first gene (No. of chromosomes examined = 52). The notation S/A, Y/F, or T/A indicates that a polymorphism was observed at that site. The notation 'not-amplified' indicates that PCR amplification failed. If the anomaloscope determined protanopia, and the amino acid combination of the first gene was A-F-A as is the second gene, indicating the absence of *OPN1LW*. On the other hand, participants determined to be deuteranopia or deuteranomaly with no increase in the second gene would reflect a lack of *OPN1MW* equivalent to trichromats. In the cases where the second gene was amplified or polymorphism was observed in male participants with deuteranopic vision, it is possible that genes coded after *OPN1MW2*, which are not expressed, were amplified. However, detailed genetic analyses, such as copy number variation and mutations in other regions, are needed to make a clear determination of genotype. Considering ethical policies, genetic information is not disclosed to participants, and age information, which can provide personal clues, is not included in the table.
